# Microglial signalling pathway deficits associated with the R47H TREM2 variant linked to AD indicate inability to activate inflammasome

**DOI:** 10.1101/2020.09.08.288043

**Authors:** Katharina Cosker, Anna Mallach, Janhavi Limaye, Thomas M Piers, James Staddon, Stephen J Neame, John Hardy, Jennifer M Pocock

**Affiliations:** Department of Neuroinflammation, University College London, Queen Square Institute of Neurology, 1 Wakefield Street, London WC1N 1PJ, UK; Eisai Ltd, Mosquito Way, Hatfield, AL10 9SN, UK; Department of Neurodegenerative Disease, UCL Queen Square Institute of Neurology, Queen Square, London WC1N 3BG, UK, Dementia Research Institute, University College, London, UCL, Reta Lila Weston Institute, UCL Queen Square Institute of Neurology, 1 Wakefield Street, London WC1N 1PJ, UK, NIHR University College London Hospitals Biomedical Research Centre and Institute for Advanced Study, The Hong Kong University of Science and Technology, Hong Kong SAR, China

**Keywords:** Alzheimer’s disease, human-iPS-microglia, NLRP3 inflammasome, R47H TREM2 variant

## Abstract

The R47H variant of the microglial membrane receptor TREM2 is linked to increased risk of late onset Alzheimer’s disease. Human induced pluripotent stem cell derived microglia (iPS-Mg) from patient iPSC lines expressing the AD-linked R47H^het^ TREM2 variant, common variant (Cv) or an R47H^hom^ CRISPR edited line and its isogeneic control, demonstrated that R47H-expressing iPS-Mg expressed a deficit in signal transduction in response to the TREM2 endogenous ligand phosphatidylserine with reduced pSYK-pERK1/2 signalling and a reduced NLRP3 inflammasome response, (including ASC speck formation, Caspase-1 activation and IL-1beta secretion). Apoptotic cell phagocytosis and soluble TREM2 shedding were unaltered, suggesting a disjoint between these pathways and the signalling cascades downstream of TREM2 in R47H-expressing iPS-Mg, whilst metabolic deficits in glycolytic capacity and maximum respiration were reversed when R47H expressing iPS-Mg were exposed to PS+ expressing cells. These findings suggest that R47H-expressing microglia are unable to respond fully to cell damage signals such as phosphatidylserine, which may contribute to the progression of neurodegeneration in late-onset AD.

## Introduction

Findings from genome-wide association studies (GWAS) point to a number of genes identified as Alzheimer’s Disease (AD) risk factors which encode for proteins expressed by microglia, the brain’s immune cells, including triggering receptor on myeloid cells 2 (TREM2) (Guerriero *et al*, 2013; Jonsson *et al*, 2013). TREM2, a transmembrane protein, functions to regulate microglial survival, proliferation and phagocytosis (reviewed in Ulland and Colonna, 2018). Whilst a number of TREM2 variants have been identified as risk factors for AD, the R47H variant, expressed in heterozygous form, confers a two-three fold increased risk of developing LOAD. The R47H variant expresses as a loss-of-function for TREM2 in mouse models (Cheng-Hathaway *et al*, 2018; Song *et al*, 2018) and impaired ligand binding (Yeh *et al*, 2016; Sudom *et al*, 2018).

Phosphatidylserine (PS), the phospholipid exposed on the external surface of damaged cells, is a ligand for TREM2 and can induce TREM2 activation in reporter cell lines (Wang *et al*, 2015;Shirotani *et al*, 2019) and phagocytosis in iPSC-derived microglia (Garcia-Reitboeck *et al*., 2018). Downstream of TREM2 activation, intracellular signalling pathways are initiated through interactions with DAP12, which upon ligand binding to TREM2 is phosphorylated and recruits Syk kinase (reviewed in Colonna and Wang, 2016; Konishi and Kiyama, 2018). Recent studies using overexpressing TREM2/DAP12 cell lines indicate that pSYK signalling induced by PS is compromised in R47H TREM2 lines (Sudom *et al*, 2018; Nugent *et al*, 2020), however pSYK signalling downstream of PS-TREM2 interactions has so far been unexplored in microglial models of TREM2 AD variants. Downstream of pSYK, pERK1/2, PI3K/pAKT and PLCγ signalling are implicated in TREM2 activity and function (Takahashi *et al*, 2005; Peng *et al*, 2010) though it is not clear whether these pathways are activated by PS and how expression of R47H TREM2 impacts signalling or cellular function in microglia.

One ongoing question is how signalling downstream of TREM2 links to inflammation in AD. TREM2 can regulate inflammatory responses (Li and Zhang, 2018) although there is conflicting evidence over whether its role is anti-inflammatory or pro-inflammatory and may depend on the cell type or the stimulus. One consequence of TREM2 signalling may involve the NLRP3 inflammasome, which is activated in microglia (Gustin *et al*, 2015) and is implicated in AD (Tan *et al*, 2013; Saresella *et al*, 2016). Aβ can induce inflammasome activation (Kono *et al*, 2014). The NLRP3 inflammasome is regulated by phosphorylation (Song and Li, 2018, Mambwe *et al*, 2019) via pSyk (Gross *et al*, 2009; Hara *et al*, 2013; Lin *et al*, 2015) and pERK1/2 signalling (Ghonime *et al*, 2014; Fann *et al*, 2018), however it is not known whether these signalling pathways downstream of TREM2 are linked to inflammasome activation.

In this study we identify functional deficits of the R47H TREM2 AD risk variant using human iPSC-derived microglia (iPS-Mg) from patients harbouring a single copy of the TREM2 R47H polymorphism (R47H^het^), in response to PS expressed on dead cells (PS+ cells) and DOPS liposomes. Here we show that iPS-Mg, expressing the TREM2 common variant (Cv), stimulated with PS+ cells show robust signalling responses that are significantly reduced in patient iPS-Mg harbouring the R47H^het^ variant, or in CRISPR edited R47H^hom^ iPS-Mg. This leads to functional deficits in NLRP3 inflammasome activation in response to the canonical LPS/ATP treatment, but also in response to PS+ cells.

## Results

### R47H^het^ patient iPS-Mg exhibit loss of signalling capacity downstream of TREM2 activation

TREM2 signalling was activated by antibody crosslinking in myeloid precursor cells and induced downstream phosphorylation events. Thus, in TREM2 common variant (Cv, control) iPS-Mg lines, activation of TREM2 by 2 min of antibody crosslinking with F(ab’)_2_ fragments resulted in increased pSYK signalling compared with IgG2B isotype control. In contrast, activation of pSYK was not observed in R47H^het^ iPS-Mg **(Fig. 1a, b)**. Additionally both pERK1/2 and pAKT signalling were activated downstream of TREM2 in Cv control iPS-Mg, (**Fig. 1a, b)** and again there was a deficit in the induction of these pathways in R47H^het^ iPS-Mg, **(Fig. 1a, b)**.

**Figure 1.**
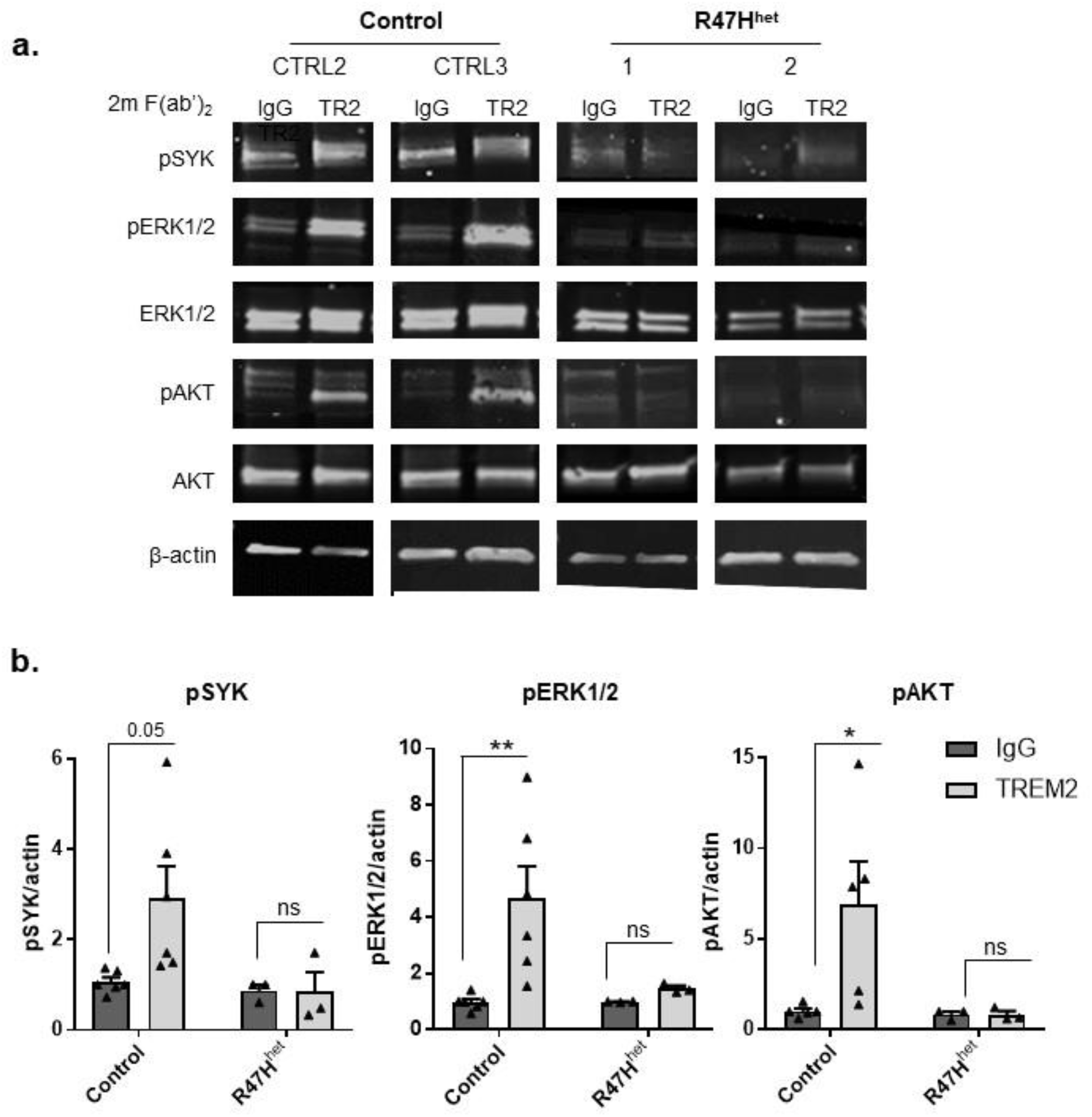
**a.** Western blot analysis of signalling pathways in iPS-derived myeloid progenitor cells following TREM2 activation with TREM2 antibody crosslinking, or IgG control, in control and TREM2 R47H^het^ patient lines. **b.** Quantification of pSYK, pERK1/2 and pAKT protein normalised to beta-actin; data are mean ± SEM, n=3-6 cell lines from 3 individual experiments, \**P*<0.05 (two-way ANOVA with Tukey’s correction), ns, non-significant.

### R47H^het^ and R47H^hom^ iPS-Mg show no deficit in phagocytosis of PS-expressing apoptotic cells or in secretion of soluble TREM2

In order to further understand the implications of an R47H signalling deficit in microglia, we cultured iPS-Mg from control and R47H TREM2^het^ lines as previously described (Xiang *et al*, 2018; Piers *et al*, 2020) and treated them with the TREM2 ligand PS, found on the outer membrane of apoptotic and necrotic cells (Nagata *et al*, 2016). The exposure of PS on the surface of heat-shocked SH-SY5Y was confirmed by Annexin-V FACS analysis **(Supplementary Fig. 1i, ii)**. We have previously shown that iPS-Mg harbouring the NHD variant T66M showed a significant deficit in phagocytosis of dead cells (Garcia-Reitboeck *et al*, 2018) and we were able to repeat this under our current conditions **(Supplementary Fig. 1iii)**. We found that in R47H^het^ iPS-Mg, there was no significant reduction in uptake of Vybrant-DiI labelled PS+ cells compared with control iPS-Mg as determined by FACS analysis **(Fig. 2a)**. We also tested this in iPS-Mg cultured from R47H^hom^ (BIONi010-C7) compared with its isogeneic control (BIONi010-C), and determined that the R47H^hom^ iPS-Mg also showed no deficit in their ability to phagocytose PS+ dead cells **(Fig. 2a, Supplementary Fig. 1iv).** We also analysed the secretion of soluble TREM2 (sTREM2) from unstimulated cells, as previously we showed that this was affected by NHD TREM2 variants expressed in iPS-Mg (Garcia-Reitboeck *et al*, 2018). Basal secretion of sTREM2 in R47H^het^ or R47H^hom^ variant iPS-Mg was not significantly different across the iPS-Mg lines or compared with control iPS-Mg lines **(Fig. 2b)**. Interestingly we found that when PS+ cells were added to iPS-Mg generated supernatant, this resulted in significantly reduced sTREM2 levels measured in the ELISA **(Supplementary Fig. 1v)** indicating that PS+ cells were able to “mop up” sTREM2. This was observed in the slight decrease in sTREM2 released following exposure of control iPS-Mg to PS+ cells, not seen in R47H^het^ or R47H^hom^ cells **(Fig. 2b).**

**Figure 2.**
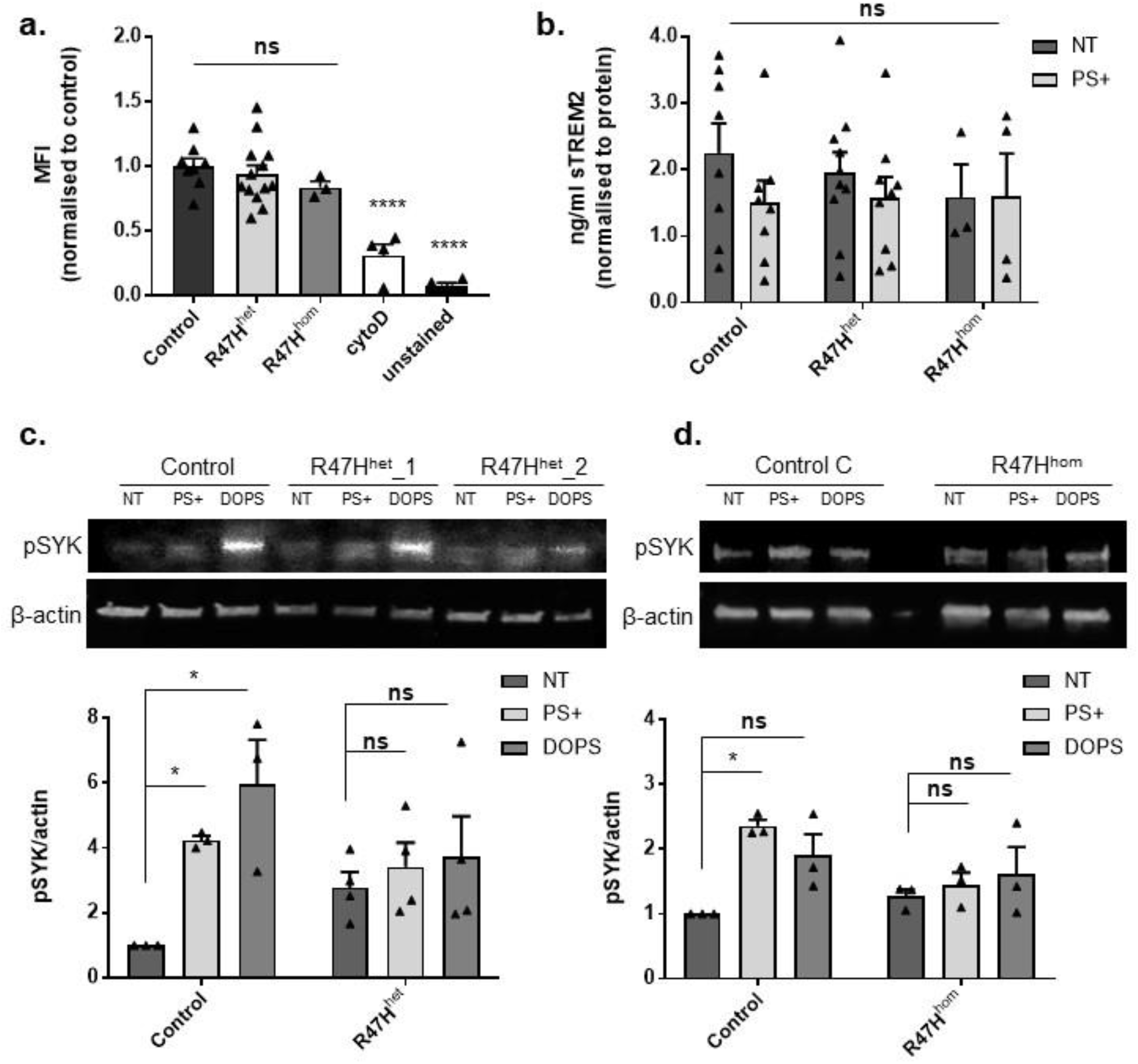
**a.** Flow cytometry of phagocytosis of DiI-labelled heat-shocked SH-SY5Y cells following 2 h iincubation with iPS-Mg in control, TREM2 R47H^het^ and R47H^hom^ lines, with cytochalasin D and unstained control; data are mean ± SEM, n=3-14 cell lines from 4 individual experiments, \*\*\*\**P*<0.0001 (one-way ANOVA with Tukey’s correction), ns =non-significant. **b**. ELISA to measure shed TREM2 in supernatant of iPS-Mg from control, TREM2 R47H^het^ and R47H^hom^ lines treated with PS+ SH-SY5Y cells; data are mean ± SEM, n=3-10 cell lines from 4 individual experiments, ns, no-significant (two-way ANOVA). **c.** Top, western blot analysis of pSYK signalling in iPS-Mg following 5 min stimulation with PS+ SH-SH5Y cells and DOPS liposomes in control and TREM2 R47H^het^ lines. Bottom, quantification of pSYK protein normalised to beta-actin; data show mean ± SEM, n=3-4 cell lines from 3 individual experiments, \**P*<0.05 (two-way ANOVA with Tukey’s correction). **d.** Top, western blot analysis of pSYK signalling in iPS-Mg following 5min stimulation with PS+ SH-SH5Y cells and DOPS liposomes in control and TREM2 R47H^hom^ lines. Bottom, quantification of pSYK protein normalised to beta-actin; data are mean ± SEM, n=3 cell lines from 3 individual experiments, \**P*<0.05 (two-way ANOVA with Tukey’s correction), ns, non-significant.

### R47H^het^ and R47H^hom^ iPS-Mg show a deficit in pSYK signalling in response to PS+ or DOPS liposomes

Since it is unknown whether an endogenous ligand for TREM2 such as PS can activate TREM2 signalling pathways in iPS-Mg, we investigated whether a 5 min stimulation with either PS+ cells or liposomes containing PS (DOPS; 1,2-dioleoyl-sn-glycero-3-phospho-L-serine) were able to induce pSYK signalling. Exposure of control iPS-Mg to PS+ cells or DOPS induced a significant increased expression of pSYK **(Fig. 2c and 2d)**. However, as with TREM2 antibody stimulation, R47H^het^ or R47H^hom^ variant iPS-Mg showed no significant increase in pSYK in response to PS+ cells **(Fig. 2c and 2d).** This was not due to any differences in the expression levels of TREM2 in the difference cell lines (Ma *et al*, 2016) **(Supplementary Fig. 1vi**).

### R47H^het^ and R47H^hom^ iPS-Mg display deficits in pERK1/2 downstream of pSYK in response to PS+ or DOPS

Since ERK1/2 and AKT are allegedly activated downstream in the canonical signalling cascade from TREM2 via SYK, (Peng *et al*, 2010), we investigated whether the deficits in pSYK signalling observed in Fig 2, were also observed for ERK and AKT signalling. We found that similar to SYK, pERK1/2 activation was reduced in R47H^het^ iPS-Mg in response to PS+ cells and DOPS liposomes **(Fig. 3a)**. This was confirmed in BIONi010-C control and R47H^hom^ isogenic cells lines **(Fig 3b)**. Phospho-AKT was also induced by PS+ cells and DOPS, however in R47H^het^ and R47H^hom^ lines, there was no significant reduction in pAKT activation **(Supplementary Fig. 2i, ii).** Therefore, we focused on pERK1/2 signalling downstream of pSYK.

**Figure 3.**
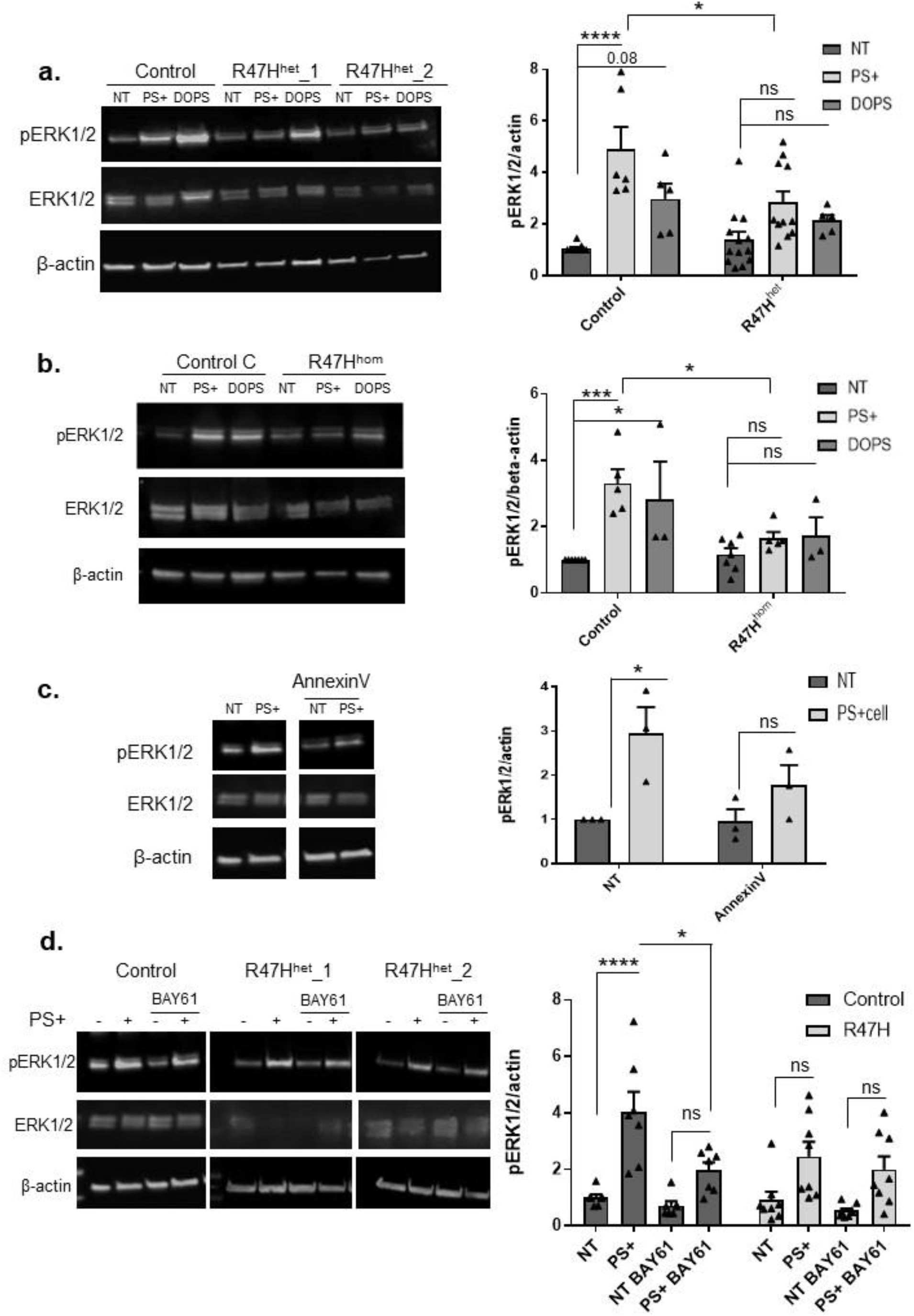
**a.** Left, western blot analysis of pERK1/2 signalling in iPS-Mg following 5 min stimulation with PS+ SH-SY5Y cells and DOPS liposomes in control and TREM2 R47H^het^ patient lines. Right, quantification of pERK1/2 protein normalised to beta-actin; data show mean ± SEM, n=5-13 cell lines from 4 individual experiments, \*\*\*\**P*<0.0001, \**P*<0.05 (two-way ANOVA with Tukey’s correction), ns, non-significant. **b.** Left, western blot analysis of pERK1/2 signalling in iPS-Mg following 5min stimulation with PS+ SH-SY5Y cells and DOPS liposomes in control and TREM2 R47H^hom^ lines. Right, quantification of pERK1/2 protein normalised to beta-actin; data show mean ± SEM, n=3-7 individual experiments, \*\*\**P*<0.001, \**P*<0.05 (two-way ANOVA with Tukey’s correction). **c.** Left, western blot analysis of pERK1/2 signalling in iPS-Mg following 5 min stimulation with PS+ SH-SY5Y cells with or without pre-incubation of recombinant Annexin-V. Right, quantification of pERK1/2 protein normalised to beta-actin; data are mean ± SEM, n=3 individual experiments, \**P*<0.05 (two-way ANOVA with Tukey’s correction). **d.** Left, western blot analysis of pERK1/2 signalling in iPS-MG following 5 min stimulation with PS+ SH-SY5Y cells with or without treatment with pSYK inhibitor 10 μM BAY61-3606 in control and TREM2 R47H^het^ patient lines. Right, quantification of pERK1/2 protein normalised to beta-actin; data show mean ± SEM, n=7-8 cell lines from 4 individual experiments, \*\*\**P*<0.0001, \**P*<0.05 (two-way ANOVA with Tukey’s correction) ns, non-significant.

We confirmed that pERK1/2 signalling activated by PS+ cells was indeed due to PS exposed on the cell surface; heat-shocked cells were pre-incubated with 5 μg recombinant Annexin-V for 1h before the PS+ cells were added to iPS-Mg. We found that Annexin-V significantly reduced the pERK1/2 response to PS+ cells, indicating that PS is responsible for activating pERK1/2 **(Fig. 3c).** To confirm that ERK phosphorylation was downstream of pSYK we used pharmacological inhibition of SYK. The specific SYK inhibitor, BAY61-3606 (10 μM) significantly inhibited the subsequent PS-induced pERK1/2 activation **(Fig. 3d)**.

### TREM2 R47H iPS-Mg are unable to activate the NLRP3-inflammasome in response to PS+ cells or LPS/ATP

SYK has been shown to be involved in activation of the NACHT-, LRR-, and pyrin (PYD)-domain-containing protein 3 (NLRP3) NLRP3-inflammasome through phosphorylation of the adaptor protein ASC (apoptosis-associated speck-like protein containing a C-terminal caspase recruitment domain) (Hara *et al*, 2013; Lin *et al*, 2015). We thus investigated whether TREM2 signalling might impact on NLRP3 inflammasome-mediated caspase-1 activation. We examined whether there were differences in ASC speck formation and subsequent caspase-1 activation in control iPS-Mg compared with R47H iPS-Mg in response to PS+ cells. As a positive control for NLRP3 inflammasome activation, cells were treated with LPS, a classical initial priming signal, and subsequently with ATP, a second activating signal for ASC-speck formation and downstream NLRP3-mediated caspase-1 activation (reviewed in Kelley *et al*, 2019). We found that significant ASC-speck formation was induced by exposure to PS+ cells or LPS/ATP in control iPS-Mg, but that R47H^Het^ or R47H^hom^ iPS-Mg showed no significant increase in ASC-speck formation above basal **(Fig. 4a)**.

**Figure 4.**
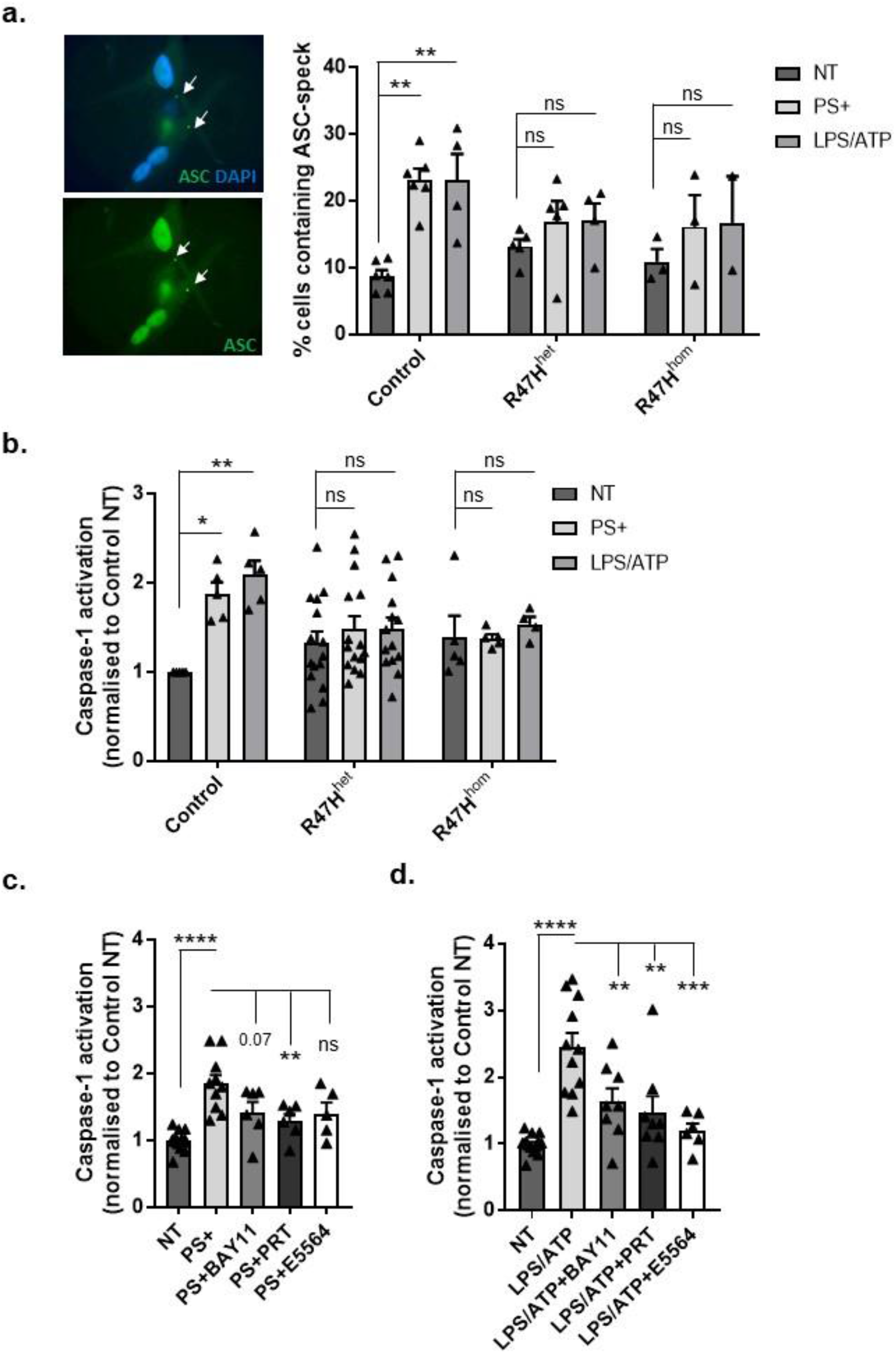
**a.** Left, immunocytochemistry of ASC specks in control iPS-Mg treated with overnight LPS + 30 min ATP, counterstained with DAPI to show cell nuclei. Right, percentage of cells containing ASC specks in iPS-Mg following treatment overnight with PS+ SH-SY5Y cells or overnight LPS + 30 min ATP in control, TREM2 R47H^het^ and R47H^hom^ lines; data are mean ± SEM, n=2-6 cell lines from 4 independent experiments, \*\**P*<0.01 (two-way ANOVA with Tukey’s correction), ns, non-significant. **b.** Caspase-1 activation in iPS-Mg following treatment overnight with PS+ SH-SY5Y cells or overnight LPS + 30 min ATP in control, TREM2 R47H^het^ and R47H^hom^ lines; data show mean ± SEM, n=5-15 cell lines from 5 individual experiments, \**P*<0.05, \*\**P*<0.01 (two-way ANOVA with Tukey’s correction). **c, d.** Caspase-1 activation in iPS-Mg treated with PS+ SH-SY5Y cells **(c.)** or LPS/ATP **(d.)** with or without inhibitors to NLRP3 (50 mM BAY11), SYK (500 nM PRT) and TLR4 (500 nM E5564); data are mean ± SEM, n=6-15 cell lines from 5 individual experiments, \**P*<0.05, \*\*\**P*<0.001 (two-way ANOVA with Tukey’s correction), ns, non-significant.

Following downstream of ASC-speck formation and activation of the NLRP3 inflammasome, inactive pro-caspase1 is subsequently cleaved to the active caspase1. Measurement of caspase1 activity with a Caspase-Glo^®^1 Inflammasome assay revealed that whilst control iPS-Mg showed significant caspase-1 activation in response to PS+ cells, R47H iPS-Mg showed a deficit in caspase 1 activation in response to PS+ cell stimulation **(Fig. 4b).** A specific caspase-1 inhibitor, YVAD-CHO was used to ensure that activation by PS+ cells was indeed attributable to Caspase-1 **(Supplementary Fig. 3i)**. We further confirmed that the activation of caspase-1 by PS+ cells was downstream of NLRP3 and SYK activation with the NLRP3 inhibitor BAY11-7082 and SYK inhibitor PRT-060318 **(Fig. 4c)**. These inhibitors also blocked caspase-1 activation by LPS/ATP treatment **(Fig. 4d)**. Pre-incubation with the TLR4 inhibitor E5564 inhibited the LPS/ATP evoked caspase activation (**Fig. 4d)**, but not the PS+ cell-induced caspase-1 response, (**Fig. 4c**), suggesting that PS+ cells signal to activate the NLRP3 inflammasome independently of TLR4 activation. In R47H^het^ and R47H^hom^ iPS-Mg, the inhibitors had no effect on caspase-1 activity, which showed no significant activation of caspase-1 with PS+ cells or LPS/ATP **(Supplementary Fig. 3ii-iv)**.

We also investigated IL-1β secretion, since the inactive precursor of IL-1β is cleaved by caspase 1 (also termed interleukin-1 converting enzyme or ICE) and subsequently secreted; a deficit was observed in the form of reduced IL-1β secretion in R47H^het^ and R47H^hom^ iPS-Mg compared with controls in response to PS+ cells (**Fig. 5a**). Interestingly stimulation with LPS/ATP evoked IL-1β release from R47H^het^ cells but not from R47H^hom^ expressing iPS-Mg although the overall level of IL-1β secretion in response to PS+ cells or LPS/ATP was less in R47H^het^ compared with Cv (**Fig. 5a**) In the canonical inflammasome pathway, LPS induces transcription of inflammasome genes, including *IL-1beta* and *NLRP3* in a priming step. Using qPCR to test whether PS+ cells also primed the iPS-Mg for NLRP3 inflammasome activation, we found that unlike LPS, neither *IL-1beta* nor *NLRP3* was transcriptionally regulated by addition of PS+ cells **(Fig. 5b, c)**. Furthermore, LPS stimulation was less effective at induction of inflammasome gene expression in R47H iPS-Mg compared with Cv **(Fig. 5b, c)**. Genes such as the alternative inflammasome *NLRP1* or *CASP1*, which has been shown not to be transcriptionally regulated, were not induced with PS+ or LPS stimulation **(Fig. 5d, e)**. We also found that *NFκB* was upregulated by LPS, but not PS+, and again this was reduced in R47H iPS-Mg compared with Cv, suggesting that transcriptional responses are affected in R47H TREM2 cells that could ultimately impact on the signalling capacity of the cell **(Fig. 5f)**.

**Figure 5.**
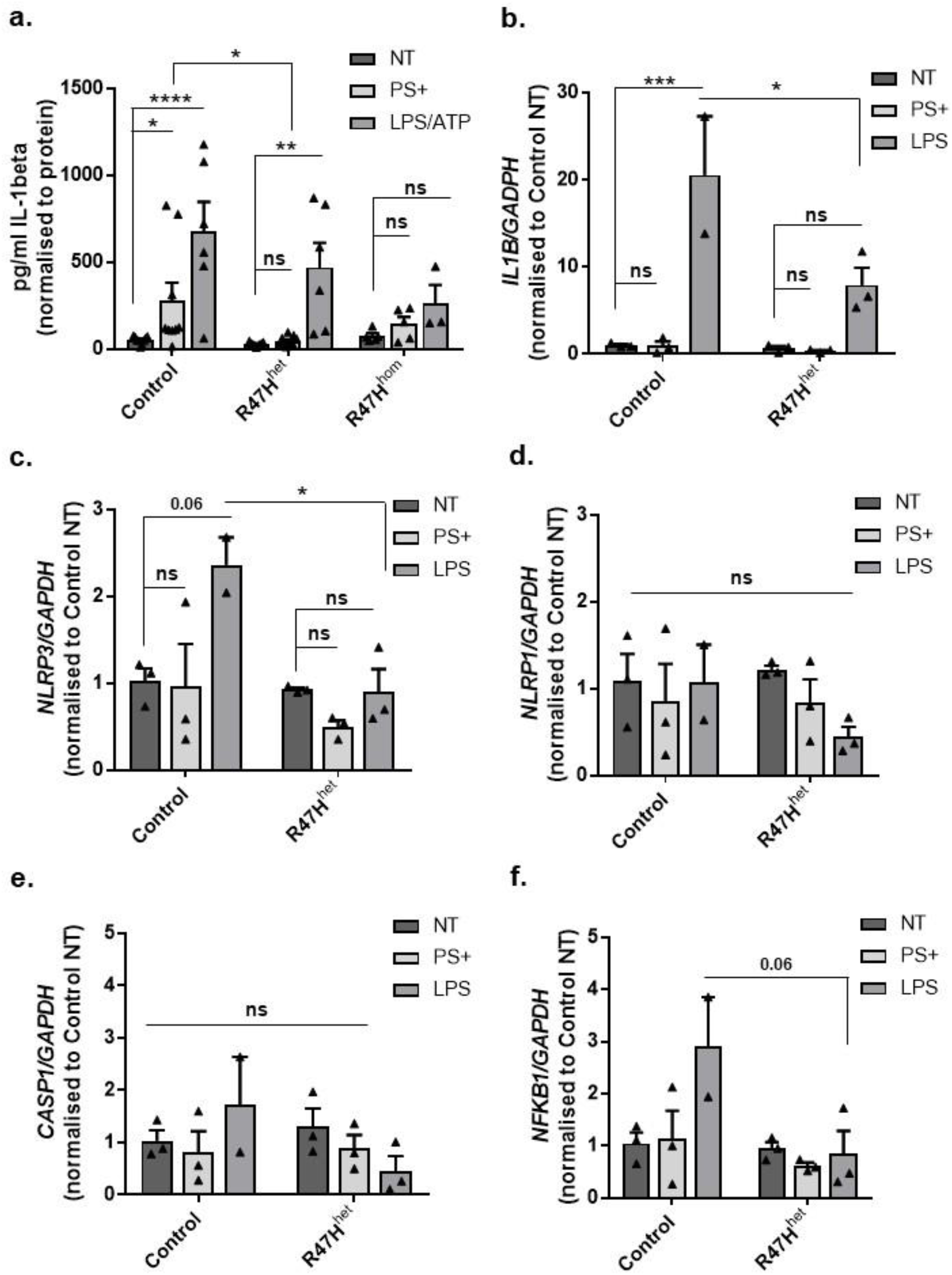
**a.** ELISA of IL-1beta protein secretion in supernatant from iPS-Mg from control, TREM2 R47H^het^ and R47H^hom^ lines following treatment overnight with PS+ SH-SY5Y cells or overnight LPS + 30 min ATP; data shown is mean ± SEM, n=3-9 cell lines from 5 individual experiments, \**P*<0.05, \*\**P*<0.01, \*\*\**P*<0.001 (two-way ANOVA with Tukey’s correction). **b.-f.** qPCR analysis in control and R47H^het^ iPS-Mg following overnight treatment with PS+ SH-SY5Y cells or LPS of **(b.)** *IL1B*, **(c.)** *NLRP3*, **(d.)** *NLRP1*, **(e.)** *CASP1* and **(f.)** *NFKB1;* data are mean ± SEM, n=2-3, \**P*<0.05, \*\*\**P*<0.001 (two-way ANOVA with Tukey’s correction), ns, non-significant.

### ECAR and OCR deficits associated with R47H are modified by PS+

Activation of the NLRP3 inflammasome is strongly linked to alterations in cell metabolism (Sanman *et al*, 2016; Wolf *et al*, 2016) and we have shown previously that TREM2 variant iPS-Mg exhibit deficits in their ability to regulate cellular metabolism (Piers *et al* 2020). In order to further understand the reduced capacity of R47H TREM2 iPS-Mg to induce an NLRP3 inflammsome response, we investigated glycolytic capability and oxidative phosphorylation by Seahorse analysis of the extracellular acidification rate (ECAR) and oxygen consumption rate (OCR), respectively, in iPS-Mg after 24 h treatment with PS+ cells. As previously reported, glycolytic and mitochondrial stress parameters revealed deficits in glycolysis and oxidative phosphorylation (**Fig. 6a, b**). Glycolytic capacity, the maximum ECAR rate, was significantly reduced in R47H^het^ iPS-Mg (**Fig. 6c**) and maximum respiration was reduced in R47H variant iPS-Mg (**Fig. 6d**). Interestingly, addition of PS+ cells rescued both the glycolytic capacity and maximum respiration in R47H^het^ iPS-Mg, but not significantly in R47H^hom^ although a similar trend was observed. These data suggest that the PS+ dead cells might act as an energy source (Xiang *et al*, 2009) that can replenish the metabolic deficiencies in R47H cells or provide lipids that feed into metabolic pathways.

**Figure 6.**
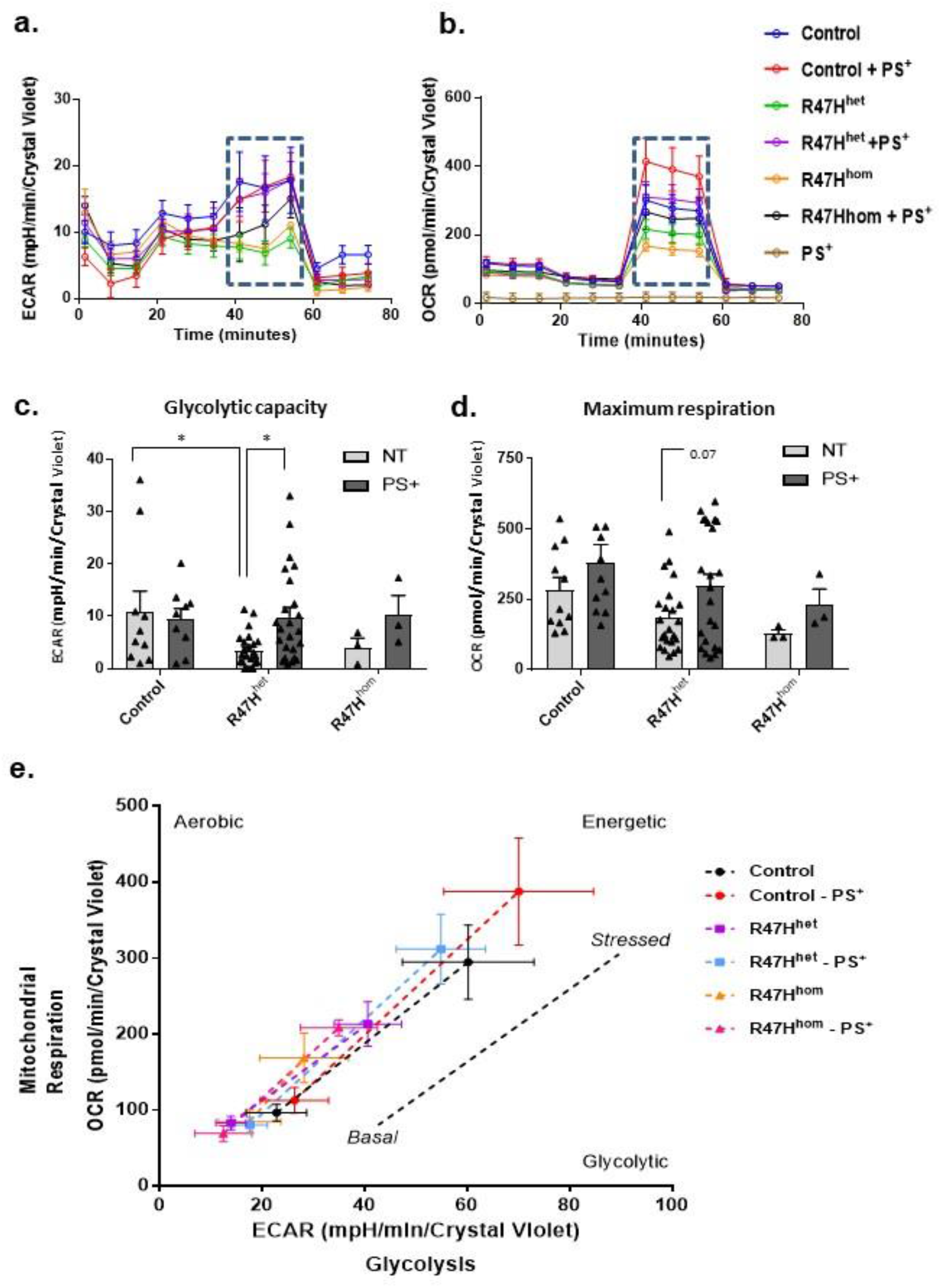
**a.** The extracellular acidification rate (ECAR) and **b.** oxygen consumption rate (OCR) were analysed in control and TREM2 R47H iPS-Mg for glycolysis and mitochondrial oxidation respectively. **c.** Analysis of iPS-Mg glycolytic capacity identified significant deficits in the ECAR of TREM2 R47H lines when compared with controls; data are mean ± SEM, n=3-24 cell lines from 3 individual experiments, \**P*<0.05 (two-way ANOVA with Tukey’s correction). **d.** Analysis of iPS-Mg maximal respiration identified reduction in the OCR of TREM2 R47H lines when compared with controls; data show mean ± SEM, n=3-23 cell lines from 3 individual experiments, \**P*=0.07 (two-way ANOVA with Tukey’s correction). **e.** Metabolic potential of iPS-Mg following FCCP treatment shows induction from a resting aerobic phenotype to an activated stressed phenotype in control and R47H TREM2 lines; data are mean ± SEM, n=3.

We previously reported that iPS-Mg expressing the R47H variant showed a reduced maximal respiratory capacity and a reduced ability to undertake a glycolytic switch when pre-incubated with an inflammatory stimulus of LPS and IFNγ (Piers *et al*, 2020). Here we found that preincubation with PS+ significantly enhanced glycolytic capacity and maximum respiration in R47H^het^ expressing iPS-Mg to similar levels found in Cv cells (**Fig 6c and d**). Glycolytic capacity and maximal respiration were not enhanced further in Cv and were not significantly rescued in R47H^hom^ expressing iPS-Mg (**Fig 6c and d**). Exposure of iPS-Mg to PS+ cells and then analysis of the cellular metabolic potential under stressful conditions (by the addition of FCCP), indicated that Cv Mg were able to increase their metabolic potential to stress when compared with CV with no PS+ cells present. The R47H variants were able to increase their metabolic potential under stressful conditions if primed with PS+ cells, but the potential was significantly below that observed in CV cells (**Fig 6e**).

## Discussion

Here we show the consequences of a reduced signalling response downstream of TREM2 in patient-derived iPS-Mg cells harbouring the R47H^het^ TREM variant, and in an R47H^hom^ line including reduced pSYK, pERK, and pAKT evident with either exposure to a PS+ ligand/DOPS liposomes or by antibody crosslinking. This reduced pSyk response is supported by findings in R47H-expressing BMDMS (Cheng-Hathaway *et al*, 2018) and R47H-expressing 293T cells (Sudom *et al*, 2018).

Whilst downstream phosphorylation events were reduced in R47H^het/hom^ iPS-Mg, phagocytosis of PS+ cells and shedding of sTREM2 at baseline were not affected. A major role for microglia in the brain is to phagocytose dead cellular material and AD is characterised by extensive neuronal loss. Whilst a role for TREM2 in phagocytosis has been proposed in a number of animal and cell models (Takahashi *et al*, 2005; Kleinberger *et al*, 2014; Wang *et al*, 2015), there is however, conflicting evidence as to whether TREM2 is involved in the phagocytosis of dead neurons; thus Takahashi *et al*, (2005) showed that shRNA knockdown of TREM2 led to reduced phagocytosis of apoptotic neurons whilst Wang *et al*, (2015) reported that microglia from Trem2-/- mice showed no difference in phagocytosis of dead cells compared with wild-type. Our findings for R47H variant iPS-Mg are different from our previous report on the more severe NHD T66M-/- patient iPS-Mg where PS+ phagocytosis was reduced (Garcia-Reitboeck *et al*, 2018), which we confirmed here. These cells do not process TREM2 to maturity, and have no TREM2 on their surface whilst R47H^het^ and R47H^hom^ iPS-Mg do express mature TREM2, and these levels were not affected by PS+ exposure (**Supplementary Fig 1 vi**). These findings are likely to underlie the differences in phagocytic responses. Phagocytosis is driven by glycolysis in macrophages (Jiang *et al*, 2016) and we reported previously that phagocytosis of Aβ was significantly affected in the R47H variant as well as T66M and W50C (Piers *et al*, 2020). However, here we find that phagocytosis of PS+ is not affected by TREM2 variants, suggesting that the effects of TREM2 variants are highly selective. Taken together, these findings would suggest furthermore that K/O models of R47H more closely resemble the T66M variant microglial phenotype than the more subtle phenotype observed in R47H^het^ or R47H^hom^ microglia which both displayed no PS+ phagocytic deficit. Additionally our current data suggest that the activation of signalling pathways through TREM2-PS interactions are independent of their ability to phagocytose dead cellular material.

We sought to determine the consequences of the downstream signalling deficit coupled to R47H. Signalling from pSyk is linked to activation of the NLRP3 inflammasome and caspase-1 (Gross *et al*, 2009; Lin *et al*, 2015). A number of different stimuli acting via specific immunoreceptor tyrosine-based activation motif (ITAMs) can induce NLRP3 inflammasome activation in microglia or macrophages (Gross *et al*, 2009; Guo *et al*, 2015; Tejera *et al*, 2019) and this includes dead cells (Kono *et al*, 2014). Here we found a reduced ability of R47H variant iPS-Mg to activate components of the NLRP3 inflammasome in response to the TREM2 ligand PS+. Thus ASC speck formation, Caspase-1 activation and IL-1β secretion, part of the canonical NLRP3 pathway (reviewed in Kelley *et al*, 2019) were reduced. Another cytokine, IL-18 is also induced upon NLRP3 activation but we were unable to detect the secretion of this cytokine (data not shown). Taken together, these findings were unexpected as TREM2 is suggested to be anti-inflammatory and thus an overall reduction in IL-1β release might be a beneficial response to dying neurons. However, transcript levels of the inflammatory cytokines IL-1β and IL-6 have been shown to be reduced in mouse models expressing mutant APP and PSEN1 in which TREM2 function was lost (Jay *et al*, 2015).

We also found that LPS/ATP was unable to induce the same degree of inflammasome response in R47H expressing microglia as Cv microglia, both at the level of priming (transcription of *IL-1β* and *NLRP3*) and ASC speck/caspase-1 /IL-1β release, suggesting a convergence in the signalling pathways of TLR4 and TREM2. Evidence of such interaction has been reported at the level of NFκB signalling which is upstream of NLRP3 activation, whilst down-regulation of TREM2 has been shown to inhibit release of inflammatory factors from LPS-stimulated microglia by inhibiting NFκB signalling pathway activity (Zhou *et al*, 2019). Interestingly *NFκB* did not show enhanced expression in Cv or R47H^het^ variant iPS microglia exposed to PS+ (**Fig. 5f**) but was significantly increased following LPS stimulation in Cv only, suggesting that *NFκB* may not be involved in PS+ stimulated pathways in human iPS Mg.

Our present findings may reflect the effect of the deficit we identified in mitochondrial respiration and glycolysis in R47H-expressing iPS-Mg. Gene transcription requires energy and microglia typically undergo a switch to glycolysis from oxidative phosphorylation to enable this to occur (Finucane *et al*, 2019), and this may explain why LPS/ATP priming is also ineffective in R47H-expressing iPS-Mg. NLRP3 inflammasome activation is linked to alterations in cell metabolism (Hughes and O’Neill 2018) and an increased glycolytic rate facilitates this activation (reviewed in Elliott and Sutterwala, 2015). NLRP3 activation can also modulate glycolysis (Finucane et al., 2019) possibly by increasing glucose uptake driven by IL-1β (Kol *et al*, 1997). We reported previously that microglia expressing the R47H variant are unable to undertake this glycolytic switch (Piers *et al*, 2020). Interestingly, despite our finding that PS+ cells exposure in the R47H expressing cells can ameliorate some of the mitochondrial deficits, the microglia still showed signalling deficits in response to PS+, suggesting that the phagocytic pathway which we found not to be affected by R47H, may be primarily responsible for this amelioration. Therefore in disease paradigms where lipid signalling is used to compensate for lack of energy, carrying an R47H variant may prevent ‘efficient’ use of the lipid energy source. Evidence suggests that TREM2 can regulate the expression of many genes influencing lipid transport such as ApoE. Our data suggest that R47H variant microglia may use PS+ cells to overcome metabolic dysfunction and in turn drive phagocytosis (reviewed in Nadjar, 2018). This may be detrimental to nearby cells transiently expressing PS on their surface as ectopic PS exposure on live neurons can cause engulfment of distal neurites (Sapar *et al*, 2018) and viable neurons (reviewed in Brown and Neher, 2014).

Taken together our findings show that R47H variant microglia express a deficit in upstream and down-stream signalling responses, culminating in their inability to activate the NLRP3 inflammasome in response to a TREM2 ligand, PS, and a reduced ability to activate the inflammasome in response to a classic inflammatory/priming signal, both of which have ramifications for microglial responses in AD.

## Materials and Methods

### Generation of Human iPSC-microglial cells (iPS-Mg)

Ethical permission for this study was obtained from the National Hospital for Neurology and Neurosurgery and the Institute of Neurology joint research ethics committee (study reference 09/H0716/64). R47H heterozygous (R47H^het^) patient-derived fibroblasts were acquired with a material transfer agreement between University College London and University of California Irvine Alzheimer’s Disease Research Center (UCI ADRC; Professor M Blurton-Jones). Control iPSC lines used in this study are as follows: CTRL1 (kindly provided by Dr Selina Wray, UCL Institute of Neurology); CTRL2 (SBAD03, Stembancc); CTRL3 (SFC840, Stembancc); CTRL4 (BIONi010-C, EBiSC). The R47H^hom^ line was a gene-edited isogenic of CTRL4 (BIONi010-C7, EBiSC). Fibroblast reprogramming, the generation of iPS-Mg and CNV analysis were performed as previously reported (Xiang et al., 2018; Piers et al., 2020). All iPSCs were maintained and routinely passaged in Essential 8 medium (Gibco). Karyotype analysis was performed by The Doctors Laboratory, London, UK (Piers et al., 2020). Using our previously described protocol, iPS-microglia (iPS-Mg) were generated (Xiang *et al*, 2018; Piers *et al*, 2020). All patient R47H lines used here are heterozygous, and thus will be referred to throughout the paper as R47H^het^ whilst the gene-edited line will be referred to as R47H^hom^.

### Antibody crosslinking experiments

iPS-derived myeloid progenitor cells were harvested and resuspended in X-VIVO medium with 100ng/ml macrophage colony stimulating factor (MCSF) at 1 million cells/ml. Myeloid progenitor cells were treated in suspension for 1 h with 5μg of IgG2B isotype control (clone 141945; R&D Systems) or αTREM2 (rat clone 237920; R&D Systems) before being treated with 10μg F(ab’)2 (Jackson #712006153) for 2 min of crosslinking. Cells were pelleted and lysed on ice in modified RIPA buffer (25 mM Tris pH7.4, 50mM NaCl, 0.5% NP-40, 0.25% Na deoxycholate, 1mM EDTA) containing 1x Halt™ protease and phosphatase inhibitor cocktail for subsequent western blot analysis.

### Apoptotic cell generation and treatment of iPS-Mg

Phosphatidylserine (PS) expression on the external surface of apoptotic SH-SY5Y cells was used as a ligand for TREM2 as we previously reported (Garcia-Reitboeck *et al*, 2018). Briefly, SH-SY5Y cells (a kind gift from Prof R. de Silva, University College London, Queen Square Institute of Neurology) were cultured in DMEM with 10% FBS (Life Technologies) and 1% penicillin/streptomycin (Life Technologies). To induce apoptosis, SH-SY5Y cells were heat shocked at 45°C for 2 h, before resuspension in iPS-Mg medium. Cell death and exposure of cell membrane PS was confirmed by flow cytometry using FITC-conjugated Annexin-V (Miltenyi Biotech) and propidium iodide staining. Apoptotic cells (“PS+ cells”) were added to iPS-Mg at a ratio of 2:1 PS+ cells:iPS-Mg for either 5 min (signalling assays) or O/N (inflammasome assays). Exposure of iPS-Mg to PS+ cells was blocked by preincubation of PS+ cells for 1 h with 5 μg recombinant Annexin V (268-10001, RayBiotech). SYK signalling was blocked by preincubation of iPS-Mg for 1 h with 10 μM BAY61-3605 prior to addition of PS+ cells.

### DOPS liposome preparation and treatment of iPS-Mg

DOPS lipids (#840035C; Avanti Polar Lipids) at 10 mg/ml in chloroform, were dried under a nitrogen gas stream in glass vials then stored under vacuum for a further 4 h prior to use. The dried lipids were hydrated with 250μl pre-chilled PBS buffer in the range of 2-10 mg/ml and sonicated in a water bath for 30 min, maintaining a temperature of ~4°C. The lipids were then processed using 21 passes through an Avanti extruder (#610000-AVL) with 100 nm polycarbonate membranes, until the solution turned from cloudy to clear. The liposomes were prepared at a 20 mM concentration in glass vials and kept at 4°C for up to 2 weeks. Functional TREM2 agonist activity was confirmed by comparing luciferase induction in GloResponse™ NFAT-RE-luc2 Jurkat Cell Line (Promega, CS176401) against the same line ectopically expressing full length human TREM2. 16-24 h before the assay Jurkat cells were plated at 5 x 105/ml in 10 % serum medium. On the assay day Jurkat cells were harvested into 1 % serum medium at 106 μl /ml and 25 μl was dispensed per well of a 384 well plate. A liposomes dilution range was dispensed at 2.5 μl per well in triplicate and luciferase activity was assessed after 16-24 h incubation. DOPS liposomes were added to iPS-Mg at 50 ng/ml for 5 min. SYK signalling was blocked by preincubation of iPS-Mg for 1 h with 10 μM BAY61-3605 prior to addition DOPS liposomes.

### Immunoblotting

Immunoblotting of Syk and ERK signalling was carried out using conventional western blotting techniques. Briefly iPS-Mg were plated at 500,000 cells/well on 6 well plates and following treatment were lysed in modified RIPA buffer (25 mM Tris pH7.4, 50 mM NaCl, 0.5% NP-40, 0.25% Na doexycholate, 1 mM EDTA) containing 1x Halt™ protease and phosphatase inhibitor cocktail (Life Technologies). Cell lysates were normalised after Bradford protein quantification and resolved on 4-20% Criterion TGX Precast Midi Protein gels (Bio-Rad), transferred to nitrocellulose and blocked in 5% milk in PBS-T (PBS + 0.05% Tween 20). Blocked membranes were incubated in primary antibody O/N at 4°C (Table 1), washed with PBS-T and incubated with corresponding LiCor compatible 680/800nm conjugated secondary antibodies (Table 1). The membranes were washed and visualised using an Odyssey detection system (LiCor). Protein bands were subsequently quantified using ImageJ software (www.imagej.nih.gov/ij).

**Table 1.**

| Antibody | Source | Cat. No. | Dilution |
| --- | --- | --- | --- |
| Phospho-AKT | Cell Signalling Technology | #4060 | 1:1000 |
| Pan-AKT | Cell Signalling Technology | #4691 | 1:1000 |
| ASC | Santa Cruz | sc-271054 | 1:100 |
| β-actin | Sigma | A5441 | 1:10000 |
| Phospho-ERK1/2 | Cell Signalling Technology | #9101 | 1:1000 |
| Pan-ERK1/2 | Cell Signalling Technology | #9102 | 1:1000 |
| Phospho-SYK/ZAP70 | Cell Signalling Technology | #2701 | 1:1000 |
| Alexa Fluor rb 790 | abcam | ab175784 | 1:10000 |
| Alexa Fluor ms 680 | Thermo Fisher | A-21058 | 1:10000 |
| Alexa Fluor ms 488 | Thermo Fisher | A-11001 | 1:500 |

### Immunostaining

Immunostaining was carried out for the localisation of ASC specks. iPS-Mg were plated at 50,000 cells/well on glass coverslips in 24 well plates and following treatment, fixed with 4% paraformaldehyde (PFA) for 20 min, followed by PBS washing, permeabilisation with 0.2% Triton for 10 min, and block (2% bovine serum albumin (BSA)) for 1 h at RT. Coverslips were incubated with primary antibody (Table 1) in blocking buffer O/N at 4°C, washed 3x in PBS and incubated with secondary antibodies (Table 1) in blocking buffer for 1 h at RT. Coverslips were washed 3x in PBS and mounted on microscope slides with Vectashield+DAPI. Images were captured using a Zeiss Axioskop 2 fluorescence microscope with 40x Neofluor objective (Oberkochen, Germany), using Axioskop software and images were analysed using ImageJ software (www.imagej.nih.gov/ij).

### Phagocytosis of apoptotic cells

SH-SY5Y cells were loaded with Vybrant CM-DiI dye (1:200; Thermo Fisher) for 15 min before heat-shock at 45°C for 2 h to induce apoptosis. Vybrant-labelled PS+ cells were added to each well of iPS-Mg (plated at 100,000 cells/well in 24-well plates) for 2 h. To block phagocytosis, iPS-Mg were incubated at 37°C +/- cytochalasin D (10 μM) for 30 min. iPS-Mg were resuspended in PBS for 10 min before FACS analysis using a Becton Dickinson FACSCalibur analyser. Data were analysed using Flowing software 2.

### Inflammasome gene array

A custom gene array for inflammasome genes was used following stimulation of iPS-Mg with PS+ cells or LPS for 24 h (TaqMan™ Array Plate 16 plus Candidate Endogenous Control Genes; Thermo Fisher Scientific). Complementary DNA was generated from iPS-Mg RNA samples using the High-Capacity RNA-cDNA kit (Life Technologies), according to the manufacturer’s instructions. Quantitative PCR was conducted on an Mx3000p qPCR system with MaxPro qPCR software (Agilent Technologies) using TaqMan™ Gene Expression Master Mix (Thermo Fisher) using a TaqMan^®^ Array Standard, 96-well Plate Format 16 + candidate endogenous controls, Catalog #: 4413265, Design ID: RAAAAE9. The following Taqman assay codes were used:

IL1B: Hs01555410_m1
NLRP1: Hs00248187_m1
NLRP3: Hs00918082_m1
CASP1: Hs00354836_m1
NFKB1:Hs00765730_m1
GAPDH: Hs99999905_m1

### Secretion of sTREM2

Supernatant was collected from iPS-Mg and centrifuged at 300 g for 15 min to remove cell debris. Soluble TREM2 (sTREM2) was measured with an in house-generated ELISA system as previously described (Garcia-Reitboeck et al., 2018). Values were normalised to protein content of cell lysates for each sample.

### Caspase-Glo^®^ 1 Inflammasome assay

Inflammasome activity was measured using Caspase-Glo^®^ 1 Inflammasome Assay (G9951; Promega). iPS-Mg were plated at 50,000 cells/well in opaque 96-well plates and treated O/N with PS+ cells. As a positive control for inflammasome activation, iPS-Mg were treated O/N with lipopolysaccharide (LPS, 100 ng/ml), + 30 min ATP (5 mM) added immediately prior to assay. iPS-Mg were preincubated for 1 h with NLRP3 inhibitor (BAY11-7082; 50 mM), pSYK inhibitor (PRT-060318; 500 nM) or the TLR4 inhibitor (E5564; 500 nM). All incubations were carried out at 37°C in the culture incubator. Cells were then allowed to equilibrate at RT for 5 min, 100 μl of Caspase-Glo^®^ 1 reagent (20 μM; Promega) was added and incubated at RT for 1 h. To verify caspase-1 specificity, a caspase-1 selective inhibitor Ac-YVAD-CHO (1 μM; Promega) was added in parallel experiments. Luminescence was measured using a Tecan plate reader.

### IL1-β ELISA

As a downstream indicator of inflammasome activation, the secretion of interleukin-1beta, (IL-1β), was analysed with a Human IL-1β/IL-1F2 R&D Quantikine ELISA Kit (#D6050). iPS-Mg were plated at 500,000 cells/well in 6 well plates and following treatment O/N with PS+ cells or O/N with LPS + 30 min ATP (5 mM), supernatant and cell lysate was collected. Quantification of IL-1β from supernatants was performed using the R&D Quantikine ELISA Kit. Human IL-1β standards and 200 μl of cell culture supernatants were added in duplicates to microplates and the ELISA was performed according to the manufacturer’s instructions. Results were normalised to total protein concentration of the corresponding cell lysate.

### Cellular respiration analysis

For the real-time analysis of extracellular acidification rates (ECAR) and oxygen consumption rates (OCR), iPS-Mg were plated and matured on Seahorse cell culture microplates and analysed using a Seahorse XFe96 Analyser (Agilent Technologies) as previously described (Piers *et al*, 2020). iPS-Mg were incubated overnight with or without PS+ cells. Glycolytic stress kits were used to analyse cellular glycolysis and Mito stress kits were used to analyse mitochondrial respiration. Data were analysed using Wave v2.4.0.6 software (Agilent Technologies).

### Statistical Analysis

Statistical analysis was carried out in GraphPad Prism using ANOVAs (one-way or two-way) with Tukey’s multiple comparison tests from at least 3 independent experiments.

## Acknowledgements

We would like to thank the patients and their families for their participation in this research project; and to Prof M. Blurton-Jones, School of Biological Sciences, University of California, Irvine, for providing us with patient fibroblasts for the R47H^het^ lines.

## Author Contributions

J. M. Pocock and K. Cosker designed the study and wrote the paper, K. Cosker carried out the majority of experiments except the following; A. Mallach analysed sTREM2 by ELISA and carried out phagocytosis assays with K. Cosker, J. Limaye performed the ASC speck analyses, T.M. Piers carried out the Seahorse analyses, and with K. Cosker the PCR analyses, S.J. Neame prepared the DOPS liposomes, S.J. Neame and J. Staddon provided scientific input to the signalling experiments and all authors appraised the final version of the paper.

---

**Supplementary Figure 1.**
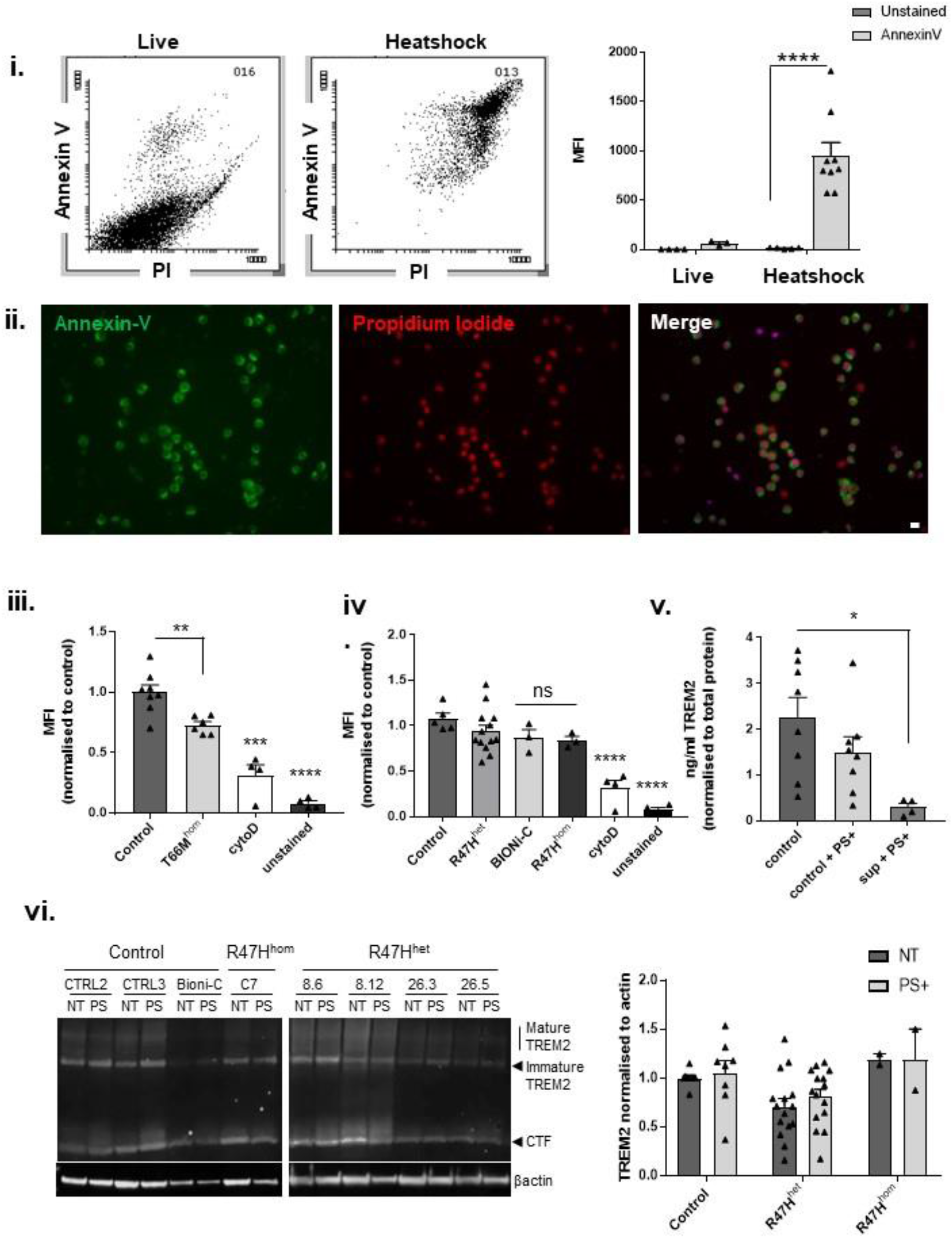
**i** Flow cytometry of Annexin-V/propidium iodide labelled SH-SY5Y cells following 2 h heat-shock at 45°C (left) and mean fluorescent intensity (right); data are mean ± SEM, n=3-9 cell lines from 3 individual experiments, \*\*\*\**P*<0.0001 (two-way ANOVA with Tukey’s correction). **ii.** Immunostaining of Annexin-V, propidium iodide and merged images of heat-shocked SH-SY5Y cells. Scale bar, 20 μm. **iii.** Flow cytometry of phagocytosis of DiI-labelled heat-shocked PS+ SH-SY5Y cells following 2 h incubation with iPS-Mg in control and T66M^hom^ lines, with cytochalasin D and unstained control; data are mean ± SEM, n=4-8 cell lines from 4 individual experiments, \*\**P*<0.01, \*\*\**P*<0.001, \*\*\*\**P*<0.0001 (one-way ANOVA with Tukey’s correction), ns, non-significant. **iv.** Flow cytometry of phagocytosis of DiI-labelled heat-shocked PS+ SH-SY5Y cells following 2 h incubation showing no difference between isogenic lines control (BIONi-C) and R47H^hom^ (BIONi-C7) lines, with cytochalasin D and unstained control; data are mean ± SEM, n=3-14 cell lines from 4 individual experiments, \*\*\*\**P*<0.0001 (one-way ANOVA with Tukey’s correction). **v.** ELISA of shed TREM2 in supernatant of iPS-Mg from control R47H^hom^ lines with or without PS+ SH-SY5Y cells treatment and supernatant incubated with PS+ SH-SY5Y cells; data are mean ± SEM, n=4-8 cell lines from 4 individual experiments, \**P*<0.05 (one-way ANOVA). **vi.** Left, western blot analysis of TREM2 expression in iPS-Mg following 5 min stimulation with PS+ SH-SH5Y cells in control and TREM2 R47H^het^ and R47H^hom^ patient lines. Right, quantification of TREM2 protein normalised to beta-actin; data are mean ± SEM, n=2-15 cell lines from 3 individual experiments, not significant (two-way ANOVA with Tukey’s correction).

**Supplementary Figure 2.**
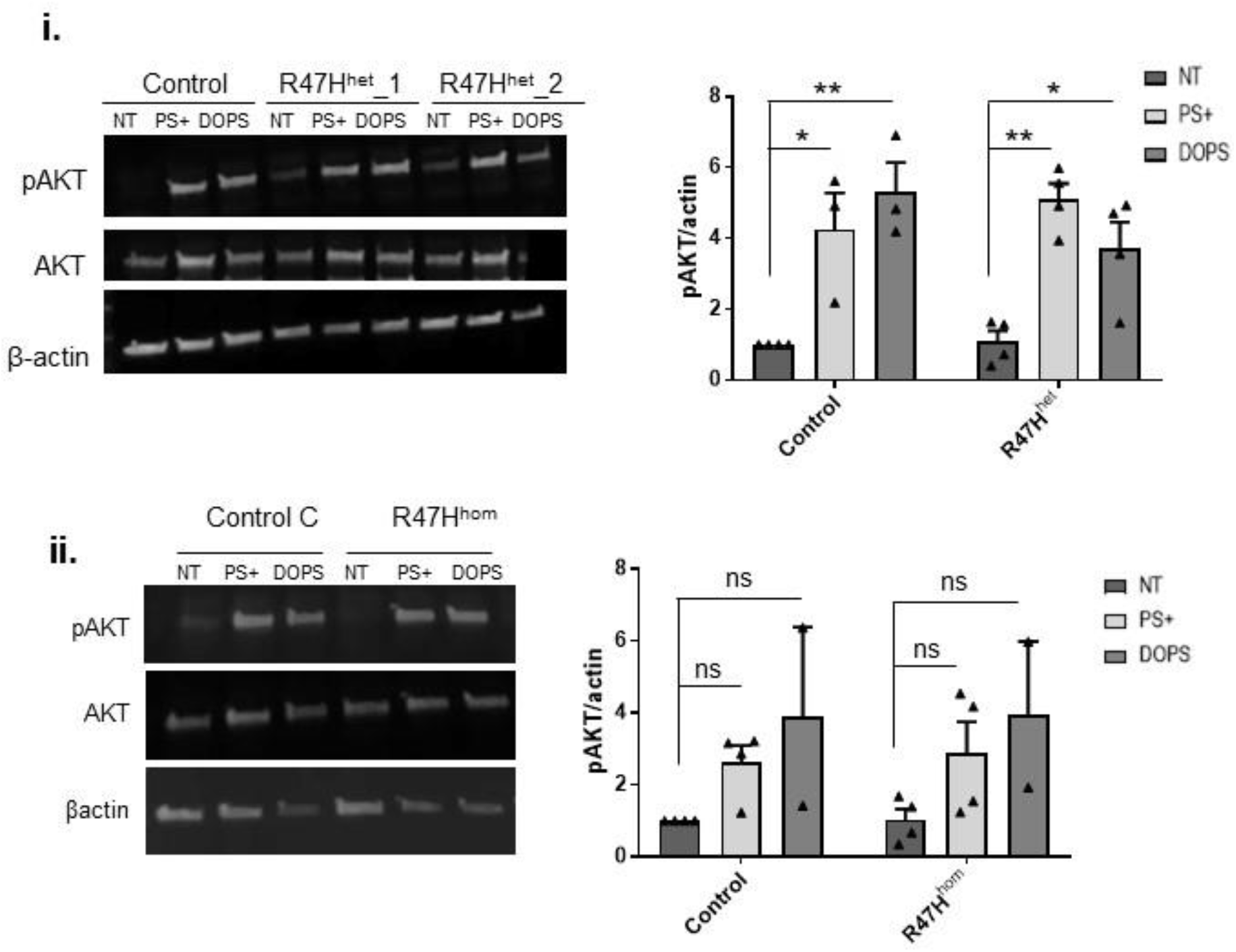
**i.** Left, western blot analysis of pAKT signalling in iPS-Mg following 5 min stimulation with PS+ SH-SY5Y cells or DOPS liposomes in control and TREM2 R47H^het^ patient lines. Right, quantification of pAKT protein normalised to ^beta^-actin; data show mean ± SEM, n=3-4 cell lines from 3 individual experiments, \**P*<0.05, \*\**P*<0.01 (two-way ANOVA with Tukey’s correction). **ii.** Left, western blot analysis of pAKT signalling in iPS-Mg following 5 min stimulation with PS+ SH-SY5Y cells or DOPS liposomes in control and TREM2 R47H^hom^ patient lines. Right, quantification of pAKT protein normalised to beta-actin; data are mean ± SEM, n=2-4 from 3 individual experiments, not significant (two-way ANOVA with Tukey’s correction).

**Supplementary Figure 3.**
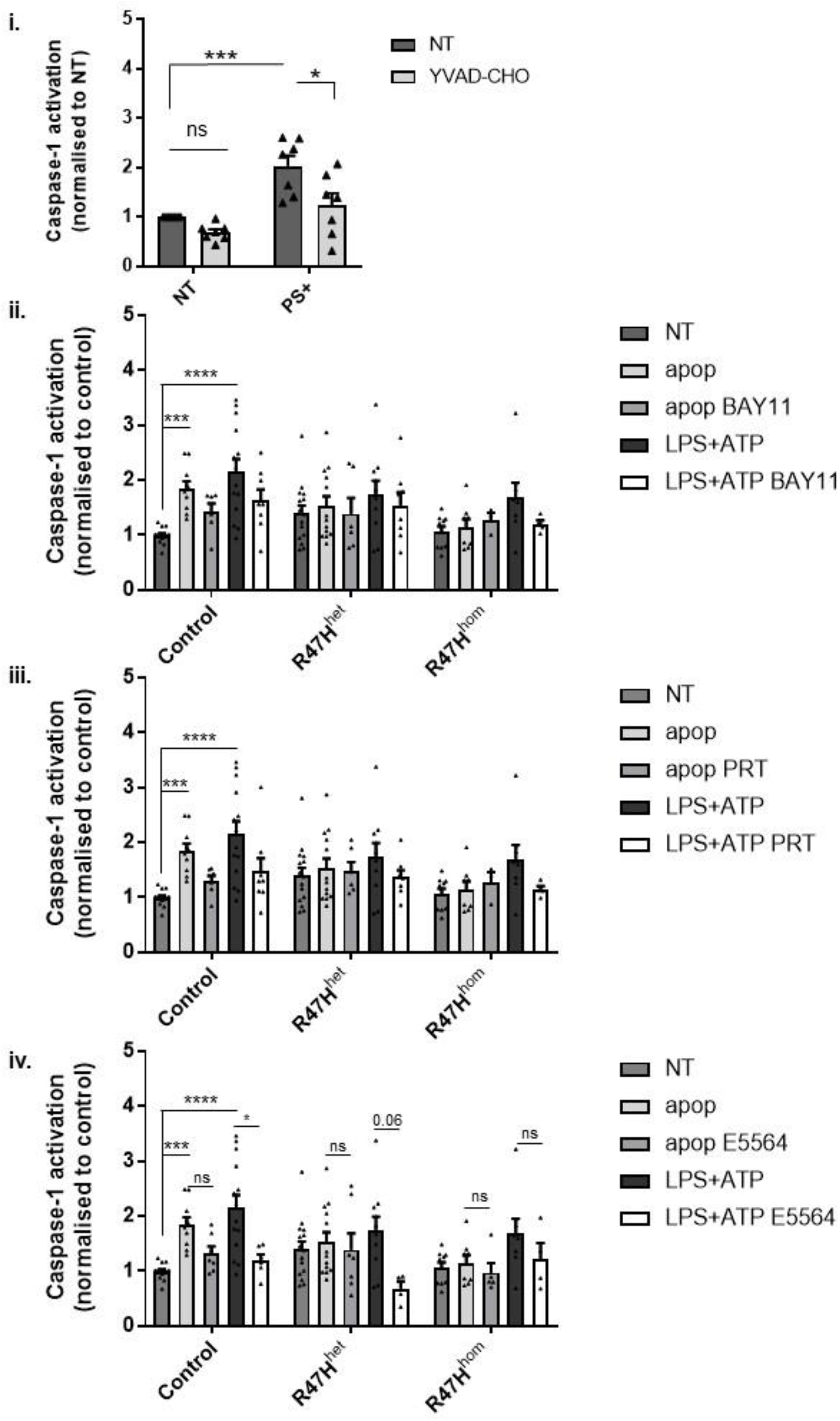
**i.** Caspase-1 activation in iPS-Mg treated with PS+ SH-SY5Y cells with or without the specific caspase-1 inhibitor YVAD-CHO; data are mean ± SEM, n=7 cell lines from 5 individual experiments, \**P*<0.05, \*\*\**P*<0.001 (two-way ANOVA with Tukey’s correction). **ii-iv.** Caspase-1 activation in iPS-Mg following treatment with **ii.,** NLRP3 (50 mM BAY11), **iii.,** SYK (500 nM PRT) or **iv.,** TLR4 (500 nM E5564) inhibitors and stimulation with overnight PS+ SH-SY5Y cells or overnight LPS + 30 min ATP in control, R47H^het^ and R47H^hom^ lines; data show mean ± SEM, n=5-15 cell lines from 4 individual experiments.

## References

Brown GC, Neher JJ, Microglial phagocytosis of live neurons. Nat Rev Neurosci 2014; 15: 209–216

Cheng-Hathaway PJ, Reed-Geaghan EG, Jay TR, Casali BR, Bemiller SM, Puntambekar SS, et al. The TREM2 R47H variant confers loss-of-function-like phenotypes in Alzheimer’s disease. Mol Neurodegen 2018; 13: 29.

Colonna M, Wang Y. TREM2 Variants: New keys to decipher Alzheimer disease pathogenesis. [Review]. Nat Rev Neurosci 2016; 17:201–7.

Elliott EI, Sutterwala FS. Initiation and perpetuation of NLRP3 inflammasome activation and assembly. Immunol Rev 2015; 265: 35–52.

Fann DYW, Lim YA, Cheng YL, Lok KZ, Chunduri P, Baik SH, et al. Evidence that NF-κB and MAPK signaling promotes NLRP inflammasome activation in Neurons Following Ischemic Stroke. Mol Neurobiol 2018; 55:1082–96.

Finucane OM, Sugrue J, Rubio-Araiz A, Guillot-Sestier M-V, Lynch MA. The NLRP3 inflammasome modulates glycolysis by increasing PFKFB3 in an IL-1β-dependent manner in macrophages. Sci Rep. 2019; 9: 4034.

Garcia-Reitboeck P, Phillips A, Piers TM, Villegas-Llerena C, Butler M, Mallach A, et al. Human induced pluripotent stem cell-derived microglia-like cells harboring TREM2 missense mutations show specific deficits in phagocytosis. Cell Rep 2018; 24:2300–11.

Ghonime MG, Shamaa OR, Das S, Eldomany RA, Fernandes-Alnemri T, Alnemri ES, et al. Inflammasome priming by lipopolysaccharide Is dependent upon ERK signaling and proteasome function. J Immunol 2014; 192:3881–8.

Gross O, Poeck H, Bscheider M, Dostert C, Hannesschläger N, Endres S, et al. Syk kinase signalling couples to the Nlrp3 inflammasome for anti-fungal host defence. Nature 2009; 459:433–6.

Guerreiro R, Wojtas A, Bras J, Carrasquillo M, Rogaeva E, Majounie E, et al. TREM2 variants in Alzheimer’s disease. N Engl J Med 2013; 368:117–27.

Gustin A, Kirchmeyer M, Koncina E, Felten P, Losciuto S, Heurtaux T, et al. NLRP3 Inflammasome Is expressed and functional in mouse brain microglia but not in astrocytes. PLoS One 2015; 10:e0130624.

Guo, H, Callaway, JB, Ting JP. Inflammasomes: Mechanism of action, role in disease, and therapeutics. Nature Medicine, 2015; 21, 677–687.

Hara H, Tsuchiya K, Kawamura I, Rang R, Hernandez-Cuellar E, Shen Y, et al. Phosphorylation of the adaptor ASC acts as a molecular switch that controls the formation of speck-like aggregates and inflammasome activity. Nat Immunol 2013; 14:1247–55.

Heneka MT, Kummer MP, Stutz A, Delekate A, Schwartz S, Saecker A, et al. NLRP3 is activated in Alzheimer’s disease and contributes to pathology in APP/PS1 mice. Nature 2013; 493:674–78.

Hughes MM, O’Neill LAJ. Metabolic regulation of NLRP3. Immunol Rev 2018; 281, 88–98.

Jay TR, Miller CM, Cheng PJ, Graham LC, Bemiller S, Broihier ML, Xu G, Margevicius D, Karlo JC, Sousa GL, Cotleur AC, Butovsky O, Bekris L, Staugaitis SM, Leverenz JB, Pimplikar SW, Landreth GE, Howell GR, Ransohoff RM, Lamb BT. TREM2 deficiency eliminates TREM2+ nflammatory macrophages and ameliorates pathology in Alzheimer’s disease mouse models. J. Exp. Med 212, 287–295 (2015).

Jonsson T, Stefansson H, Steinberg S, Jonsdottir I, Jonsson PV, Snaedal J, et al. Variant of TREM2 associated with the risk of Alzheimer’s disease. N Engl J Med 2013; 368:107–16.

Kelley N, Jeltema D, Duan Y, He Y. The NLRP3 inflammasome: An overview of mechanisms of activation and regulation. [Review]. Int J Mol Sci 2019; 20:3328.

Kleinberger G, Yamanishi Y, Suárez-Calvet M, Czirr E, Lohmann E, Cuyvers E, Struyfs H, Pettkus N, Wenninger-Weinzierl S et al. TREM2 mutations implicated in neurodegeneration impair cell surface transport and phagocytosis. Sci Transl Med. 2014; 6:243–286.

Kol S, Ben-Shlomo I, Ruutiainen K, Ando M, Davies-Hill TM, Rohan RM, Simpson IA, Adashi EY. The midcycle increase in ovarian glucose uptake is associated with enhanced expression of glucose transporter 3. Possible role for interleukin-1, a putative intermediary in the ovulatory process. J Clin Invest 1997; 99:2274–83.

Konishi H, Kiyama H. Microglial TREM2/DAP12 Signaling: A Double-Edged Sword in Neural Diseases. [Review]. Front Cell Neurosci 2018; 12:206.

Kono H, Kimura Y, Latz E. Inflammasome Activation in Response to Dead Cells and Their Metabolites. [Revew]. Curr Opin Immunol 2014; 30:91–8.

Li JT, Zhang Y. TREM2 regulates innate immunity in Alzheimer’s disease. [Review]. J Neuroinflammation 2018; 15:107.

Lin YC, Huang DY, Wang JS, Lin YL, Hsieh SL, Huang KC, et al. Syk is Involved in NLRP3 inflammasome-mediated caspase-1 activation through adaptor ASC phosphorylation and enhanced oligomerization. J Leukoc Biol 2015; 97:825–35.

Ma L, Allen M, Sakae N, Ertekin-Taner N, Graff-Radford NR, Dickson DW, et al. Expression and processing analyses of wild type and p.R47H TREM2 variant in Alzheimer’s disease brains. Mol Neurodegener 2016; 11:72.

Mambwe B, Neo K, Khameneh HJ, Leong KWK, Colantuoni M, Vacca M, et al. Tyrosine Dephosphorylation of ASC Modulates the Activation of the NLRP3 and AIM2 Inflammasomes. Front Immunol 2019; 10:1556.

Nagata S, Suzuki J, Segawa K, Fugii T. Exposure of Phosphatidylserine on the Cell Surface. [Review]. Cell Death Differ 2016; 23:952–961.

Nadjar A. Role of metabolic programming in the modulation of microglia phagocytosis by lipids. Review; Prostaglandins Leukot Essent Fatty Acids. 2018; 135:63–73.

Nugent AA, Lin K, van Lengerich B, Lianoglou S, Przybyla L, Davis SS, Llapashtica C, Wang J, TREM2 Regulates Microglial Cholesterol Metabolism upon Chronic Phagocytic Challenge. Neuron 2020; 105:837–854.

Peng Q, Malhotra S, Torchia JA, Kerr WG, Coggeshall KM, Humphrey MB. TREM2- And DAP12-dependent Activation of PI3K Requires DAP10 and Is Inhibited by SHIP1. Sci Signal 2010; 3:38.

Piers TM, Cosker K, Mallach A, Johnson GT, Guerreiro R, Hardy J, Pocock JM. A Locked Immunometabolic Switch Underlies TREM2 R47H Loss of Function in Human iPSC-derived Microglia. FASEB J 2020; 34:2436–50.

Sanman LE, Qian Y, Eisele NA, Ng TM, van der Linden WA, Monack DM, Weerapana E, Bogyo M. Disruption of glycolytic flux is a signal for inflammasome signaling and pyroptotic cell death. Elife 2016 5:e13663.

Sapar ML, Ji, H, Wang B, Poe AR, Dubey K, Ren X, Ni J-Q, Han C. Phosphatidylserine Externalization Results from and Causes Neurite Degeneration in Drosophila. Cell Reports 2018; 24: 2273–2286.

Saresella M, La Rosa F, Piancone F, Zoppis M, Marventano I, Calabrese E, et al. The NLRP3 and NLRP1 inflammasomes are activated in Alzheimer’s disease. [Review]. Mol Neurodegen 2016; 11:23.

Shirotani K, Hori Y, Yoshizaki R, Higuchi E, Colonna M, Saito T, et al. Aminophospholipids Are Signal-Transducing TREM2 Ligands on Apoptotic Cells. Sci Rep 2019; 9:7508.

Song WM, Joshita S, Zhou Y, Ulland TK, Gilfillan S, Colonna M. Humanized TREM2 Mice Reveal Microglia-Intrinsic and -Extrinsic Effects of R47H Polymorphism. J Exp Med 2018; 215:745–60.

Song N, Li T. Regulation of NLRP3 Inflammasome by Phosphorylation. [Review]. Front Immunol 2018; 9:2305.

Sudom A, Talreja S, Danao J, Bragg E, Kegel R, Min X, et al. Molecular Basis for the Loss-Of-Function Effects of the Alzheimer’s Disease-Associated R47H Variant of the Immune Receptor TREM2. J Biol Chem 2018; 293:12634–46.

Tan MS, Yu JT, Jiang T, Zhu XC, Tan L. The NLRP3 Inflammasome in Alzheimer’s Disease. [Review]. Mol Neuro 2013; 48:875–82.

Takahashi K, Rochford CDP, Neumann H. Clearance of apoptotic neurons without inflammation by microglial Triggering receptor expressed on myeloid cells-2. J Exp Med 2005; 201:647–57.

Tejera D, Mercan D, Sanchez-Caro JM, Hanan M, Greenberg D, Soreq H, Latz E, Golenbock D, Heneka MT. Systemic inflammation impairs microglial Aβ clearance through NLRP3 inflammasome. EMBO J. 2019; 2; 38(17)

Turnbull IR, Gilfillan S, Cella M, Aoshi T, Miller M, Piccio L, Hernandez M, Colonna M. Cutting edge: TREM-2 attenuates macrophage activation. J Immunol. 2006; 177:3520–3524.

Ulland TK, Colonna M. TREM2 – A Key Player in Microglial Biology and Alzheimer Disease. [Review]. Nat Rev Neurol 2018; 14:667–75.

Wang Y, Cella M, Mallinson K, Ulrich JD, Young KL, Robinette ML, et al. TREM2 Lipid Sensing Sustains the Microglial Response in an Alzheimer’s Disease Model. Cell 2015; 160:1061–71.

Wolf AJ, Reyes CN, Liang W, Becker C, Shimada K, Wheeler ML, Cho HC, Popescu NI, Coggeshall KM, Arditi M, Underhill DM. Hexokinase Is an Innate Immune Receptor for the Detection of Bacterial Peptidoglycan. Cell. 2016; 166:624–636.

Xiang X, Piers TM, Wefers B, Zhu K, Mallach A, Brunner B, et al. The Trem2 R47H Alzheimer’s Risk Variant Impairs Splicing and Reduces Trem2 mRNA and Protein in Mice but Not in Humans. Mol Neurodegener 2018; 13:49.

Yeh FL, Wang Y, Tom I, Gonzalez LC, Sheng M. TREM2 Binds to Apolipoproteins, Including APOE and CLU/APOJ, and Thereby Facilitates Uptake of Amyloid-Beta by Microglia. Neuron 2016; 91:328–40.

Zhou J, Yu W, Zhang M, Tian X, Li Y, Lü Y. Imbalance of Microglial TLR4/TREM2 in LPS-Treated APP/PS1 Transgenic Mice: A Potential Link Between Alzheimer’s Disease and Systemic Inflammation Neurochem Res 2019;44:1138–1151.

